# A sensation for inflation: initial swim bladder inflation in larval zebrafish is mediated by the mechanosensory lateral line

**DOI:** 10.1101/2023.01.12.523756

**Authors:** Alexandra Venuto, Stacey Thibodeau-Beganny, Josef G. Trapani, Timothy Erickson

**Affiliations:** Department of Biology, East Carolina University, Greenville, NC, USA; Department of Biology and Neuroscience Program, Amherst College, Amherst, MA, USA; Department of Biology, University of New Brunswick, Fredericton, NB, Canada

**Keywords:** behavior, buoyancy, hair cells, hydrodynamics, lateral line, *lhfpl5*, swim bladder, sensory system, zebrafish

## Abstract

Larval zebrafish achieve neutral buoyancy by swimming up to the surface and taking in air through their mouths to inflate their swim bladders. We define this behavior as ‘surfacing’. Little is known about the sensory basis for this underappreciated behavior of larval fish. A strong candidate is the mechanosensory lateral line, a hair cell-based sensory system that detects hydrodynamic information from sources like water currents, predators, prey, and surface waves. However, a role for the lateral line in mediating initial inflation of the swim bladder has not been reported.

To explore the connection between the lateral line and surfacing, we utilized a genetic mutant (*lhfpl5b^-/-^*) that renders the zebrafish lateral line insensitive to mechanical stimuli. We observe that approximately half of these lateral line mutants over-inflate their swim bladders during initial inflation and become positively buoyant. Thus, we hypothesize that larval zebrafish use their lateral line to moderate interactions with the air-water interface during surfacing to regulate swim bladder inflation. To test the hypothesis that lateral line defects are responsible for swim bladder over-inflation, we show exogenous air is required for the hyperinflation phenotype and transgenic rescue of hair cell function restores normal inflation. We also find that chemical ablation of anterior lateral line hair cells in wild type larvae causes hyperinflation. Furthermore, we show that manipulation of lateral line sensory information results in abnormal inflation. Finally, we report spatial and temporal differences in the surfacing behavior between wild type and lateral line mutant larvae. In summary, we propose a novel sensory basis for achieving neutral buoyancy where larval zebrafish use their lateral line to sense the air-water interface and regulate initial swim bladder inflation.

## Introduction

The swim bladder of teleost fish provides for neutral buoyancy in the water column, thereby minimizing the energy expenditure associated with locomotion (Alexander, 1966). Larval teleosts initate swim bladder inflation by swimming up to the surface to ingest air using a set of behaviors we refer to as “surfacing” (Evans and Damant, 1928; Lindsey et al., 2010; von Ledebur, 1928). Although surfacing poses a significant risk of predation, larval fish are highly motivated to reach the surface to intake air. For example, pre-inflated lake trout larvae will swim hundreds of feet at a constant rate in attempts to access the surface (Tait, 1960). Surfacing is a complex task that requires a combination of newly developed physical features, sensory systems, and motor skills (Bailey and Doroshov, 1995; Chatain, 1989; Doroshev et al., 1981; Kimmel et al., 1995; Kitajima et al., 1994) – (i) Larvae must discriminate up from down and (ii) be able to move directionally towards the surface; (iii) they must appropriately sense the air-water interface, and (iv) be able to intake air through the mouth upon arrival at the surface. Ingested air is then moved by peristalsis into the swim bladder through the pneumatic duct that connects the gut and swim bladder (Doroshev et al., 1981; Rieger and Summerfelt, 1998; Tait, 1960). Together, surfacing represents perhaps the most complex behavior that larvae perform prior to achieving neutral buoyancy (Fero et al., 2011).

Little is known about the sensory cues that instruct the surfacing behavior. Perturbations to the vestibular system typically result in hypoinflation (Nicolson et al., 1998; Schoppik et al., 2017). The inability to detect gravity by the utricular otolith is specifically responsible for the phenotype, presumably because larvae are unable to orient their movements upwards towards the surface (Riley and Moorman, 2000). Photosensory cues may also play a role in directional orientation and surface detection. However, which photosensory systems are involved has not been defined and the effects of altering lighting conditions varies between species (Stuart and Drawbridge, 2012; Trotter et al., 2003; Villamizar et al., 2009). Even less is known about how larval fish detect the air-water interface itself. Fish and some amphibians possess an additional hair cell-based sensory system-the *mechanosensory lateral line*-that contributes to detection of near-field hydrodynamic information originating from water currents, predators and prey, conspecifics, abiotic objects, and surface waves (Anneser et al., 2020; Bleckmann, 2006; Butler and Maruska, 2016; Carrillo and McHenry, 2016; Mogdans, 2019; Montgomery et al., 1997; Stewart et al., 2013; Venuto et al., 2022). Specific to surface detection, surface-feeding fish use their lateral line to detect surface waves created by their prey (Bleckmann and Schwartz, 1982). Furthermore, Japanese flying fish are predicted to use their ventrally-located lateral line to sense transitions through the air-water interface (Gibbs, 2004; Tsukamoto and Yoshino, 1957). While the lateral line is a strong candidate to aid in surfacing behaviors that lead to initial swim bladder inflation in larval fish, its specific role in this important behavior has not been explored.

A CRISPR-Cas9 knockout of *LHFPL tetraspan subfamily member 5b* (*lhfpl5b*) is the first genetic zebrafish mutant where the lateral line is non-functional from birth but hearing and balance are normal (Erickson et al., 2020). These mutants provide an opportunity to uncover novel roles for the lateral line. Unexpectedly, homozygous mutant larvae present with a hyperinflation phenotype characterized by over-filled swim bladders and positive buoyancy, a very rare finding among swim bladder defects (Abbas and Whitfield, 2009; McCune and Carlson, 2004). Using several genetic and chemical techniques to manipulate lateral line function, we show that lateral line sensory information is used by larval zebrafish to appropriately fill their swim bladders, likely by detecting surface tension at the air-water interface using mechanosensory neuromasts located on their heads. Overall, our study uncovers a novel role for the mechanosensory lateral line mediating initial inflation of the swim bladder.

## Methods

### Animal husbandry and ethics statement

Adult zebrafish (*Danio rerio*) were maintained and bred using standard procedures (Westerfield, 2000). All experiments used larvae at 2-6 days post-fertilization (dpf), which are of indeterminate sex at this stage. Except where otherwise stated, animals were placed in clear, plastic tubs (dimensions: 15 cm x 10 cm x 4 cm, E3 volume: 460 mL) at 2 dpf and maintained at 28.5°C on a 14:10 light/dark cycle at 500-700 lux in E3 embryo media (5 mM NaCl, 0.17 mM KCl, 0.33 mM CaCl2, 0.33 mM MgCl2, buffered with NaHCO3). Unhatched embryos were manually dechorionated before placing larvae in experimental conditions at 2 dpf. It is worth noting that there was no obvious difference in wild type versus *lhfpl5b* mutant hatching rate, though this was not explicitly studied. Animal research complied with guidelines stipulated by the Institutional Animal Care and Use Committees at East Carolina University (protocol D372) and Amherst College (assurance number 3925-1 with the Office of Laboratory Animal Welfare).

### Mutant and transgenic fish lines

The zebrafish mutant allele *lhfpl5b^vo35^* and transgenic lines *Tg(myo6b:EGFP-lhfpl5a)vo23* and *Tg(myo6b:EGFP-PA)vo68* were used in this study (Erickson et al., 2017; Erickson et al., 2020). For all experiments, *lhfpl5b^vo35^*homozygotes were identified from their hetero-and homozygous wild type siblings by lack of FM 1-43 (ThermoFisher) labeling of the lateral line hair cells.

To create a fish line that expresses the Channelrhodopsin-2 (ChR2) optogenetic protein solely in the hair cells of the lateral line, approximately 2 kb of DNA upstream of the start codon of the *lhfpl5b* gene was cloned into the p5E plasmid of the recombination-based multisite Gateway cloning system (ThermoFisher). A Tol2 transposon backbone was then used to create the injectable expression plasmid *lhfpl5b2028:ChR2-EYFP-PA*. This plasmid was then co-injected with transposase mRNA into single-cell embryos (Kwan et al., 2007) to create the *Tg(lhfpl5b2028:ChR2-EYFP-PA)* zebrafish line.

### Buoyancy tests

To ascertain the buoyancy status of larval fish, larvae were anesthetized with MS-222 (Western Chemical Inc., Ferndale, WA) and placed at the vertical midway point of a 50 mL conical tube containing E3 media. At the expiration of a 30-second timer, the end position of the fish was recorded and the fish was removed from the tube for imaging under the microscope. If the fish reached the top or bottom of the tube before 30 seconds, then the time it took to reach that location was recorded. The total height of the water column was 8.9 cm, and the fish was placed half-way at 4.45 cm. The rate of buoyancy was determined by dividing the centimeters travelled by the seconds it took to travel that distance (cm/s). Over-inflated fish were categorized by larvae floating to the top of the tube in 30 seconds, while under-inflated fish sunk to the bottom of the tube. Neutrally boutant or regular-inflated larvae remained near the middle of the water column. This test was conducted during the imaging portion at the end of each experiment, with the exception of the oil-water interface experiment where buoyancy could not be accurately determined due to the greater density of oil compared to air.

### Imaging and swim bladder measurements

Swim bladders were imaged using a SteREO Discovery.V8 microscope (Zeiss) equipped with a Gryphax Arktur camera (Jenoptik). Lateral and dorsal images of the swim bladder were taken for each fish. Swim bladder measurements were conducted using the measurement tool in Adobe Photoshop where measurements were calibrated according to the image of the stage micrometer taken when imaging experiments. The major axis and minor axis of the lateral view of the swim bladder was measured as well as the minor axis of the dorsal view. All measurements were taken from numbered images so that the measurer was blinded to the genotype/condition. These meaurements were halved and used in the equation for the volume of an ellipsoid: *V* = 4/3π*abc* to find the volume of the swim bladder (Lindsey et al., 2010).

### Manipulation of photosensory cues during surfacing

At 0 dpf, zebrafish embryos were collected and immediately placed in plastic tubs filled with E3 (detailed above) in an unlit incubator. Larvae were not brought into light until 6 dpf, at which point lateral line mutants and wild type siblings were identified by FM 1-43 labeling. After sorting, larvae were analyzed for buoyancy. The experiment was repeated three times using 20-21 larvae (combined mutant and wild type) per trial.

### Blocking access to the air-water interface

Mutant larvae and wild type siblings were sorted at 2 dpf into glass cylinders (diameter: 9.5 cm, E3 volume: 425 mL) equipped with fitted, wire mesh filter plungers (Bodum Inc., USA). Conditions were as follows: wild type with open access to the surface, wild type with blocked access to the surface, mutant with open access to the surface, and mutant with blocked access to the surface. All containers were filled half way with E3 and in the experimental condition (“blocked access”), the filter was pressed down below the surface for both wild type and mutant larvae. The control condition used the same amount of E3 but the filter remained above the surface so that larvae could access the exogenous air. Conditions remained constant until 6 dpf when larvae were imaged and analyzed for swim bladder inflation. The experiment was repeated three times using 15-25 larvae per condition.

### Analysis of surfacing behavior

Wild type and *lhfpl5b^vo35^* larvae were were sorted at 2 dpf and blocked from having access to the surface until 9 AM on 4 dpf when individuals were immediately placed into clear rectangular containers (dimensions: 3 cm x 3 cm x 3 cm, E3 volume: 20 mL) for video recording. Recordings lasted 12 hours, from 9 AM to 9 PM on 4 dpf. At the end of filming, larvae were imaged and analyzed for buoyancy and swim bladder volume, as described. All surfacing videos were numbered for blind analysis. Surfacing analysis was conducted by manual observation of each individual larvae from the videos (30 frames per second) and manually counting visits made to the surface (counted as reaching the meniscus), the timeframe of the video at which the surfacing event occurred, and the duration of time spent at the surface per surfacing event. Recording was repeated six times with both mutant wild type larvae present in each recording. Larvae that did not inflate their swim bladders by the end of filming (5 total larvae, 1 wild type and 4 mutants) were excluded from this analysis due to lack of surfacing events.

### Lateral line hair cell ablations

For all experiments, wild type and *lhfpl5b* mutant larvae were treated with ototoxin with untreated siblings acting as controls (n = 7-15 larvae per condition per trial, 4 trials) For single treatments, surface accessed was blocked as described above for untreated and treated larvae until 9 AM on 4 dpf when treatment occurred. Single 30 minute treatments with neomycin sulfate (EMD Millipore Corp., USA) were done with a 100 µm solution in E3 while repeated neomycin treatments (Venuto and Erickson, 2021) were done as previously described with a concentration of 50 µM. For full ablation with copper sulfate (CuSO4; Ward’s Science, Rochester, NY), the treatment concentration was 10 uM with a 30 minute exposure. For experiments where head and trunk neuromasts were selectively ablated with CuSO4, anesthetized larvae were moved to a rectangular plate (4.5 cm x 3 cm x 1 cm) coated with Sylgard (Electron Microscopy Sciences, Hatfield, PA). After removing excess water, a thin layer of petroleum jelly (Vaseline, Unilever USA) was placed horizontally at the back of the head. Next, 30 uM CuSO4 or E3 was pipetted onto the appropriate region depending on the experimental condition. Immediately following the 13 minute treatment, larvae were placed in fresh E3 in clear tubs detailed above for 24 hours until 9 AM on 5 dpf when larvae were imaged and analyzed for swim bladder inflation. For all experiments, larvae were imaged and analyzed for swim bladder inflation at 9 AM on 5 dpf. Each experiment was repeated three times using 10-24 larvae per condition.

### Transgenic rescue of lateral line mutant

Larval zebrafish from crosses of *lhfpl5b^+/-^* and *Tg(myo6b:EGFP-lhfpl5a)vo23Tg; lhfpl5b^+/-^*were sorted at 2 dpf for transgene expression. At 6 dpf, larvae were numbered, imaged for swim bladder inflation and genotyped for the *lhfpl5b^vo35^*allele as previously described (Erickson et al., 2020). The experiment was repeated twice using 26-48 larvae per condition.

### Optogenetic activation of lateral line hair cells with Channelrhodopsin-2 (ChR2)

After blocking access to the surface until 4 dpf, we exposed wild type and lateral line mutants, with or without the *lhfpl5b2028:ChR2-EYFP-PA* transgene, to LED blue light (470 nm) with an intensity of 730 lux over their holding tanks against a dark background. Light flashed for 25 milliseconds at one second intervals. Control larvae were kept in the same background, without blue light exposure. Conditions were held constant until 6 dpf when larvae were sorted for genotype, imaged and analyzed for swim bladder inflation. The experiment was repeated three times using 15-25 larvae per condition.

### Oil-water interface experiments

Mutant larvae and wild type siblings were sorted at 2 dpf into clear cylindrical containers (diameter: 5 cm, volume: 60 mL) with embryo media only or embryo media with a 2 mm thick surface layer of lab grade mineral oil (Ward’s Science, Rochester, NY). Oil on the surface decreases surface tension (Johansen, 1924) and has been previously used in experiments involving the zebrafish swim bladder (Ehrlich and Schoppik, 2017). Conditions remained constant until 6 dpf when larvae were imaged and analyzed for swim bladder volume. Because oil-filled larvae are negatively buoyant, we did not perform buoyancy tests and classified larvae as over-filled based on swim bladder volume alone. The experiment was repeated three times using 15-25 larvae per condition.

### Graphs and statistical tests

All graphs and statistical tests were done using R (R Core Team, 2022). One-way ANOVAs were used to compare the conditions as a whole (genotype independent variable plus environment independent variable) with Tukey post-hoc test. Chi-squared tests were used to analyze proportion data for over and under-inflation. For the over-inflation comparisons, the two categories used in the comparison between conditions/genotypes were (i) over-inflation and (ii) all other inflation (regular and under), meaning that totals per category/genotype accounted for 100% of the population. Similarly, for the under-inflation comparisons, the two categories used in the comparison between conditions/genotypes were (i) under-inflation and (ii) all other inflation (regular and over). P-values less than 0.05 were considered significant.

## Results

### Lateral line mutants exhibit a swim bladder hyperinflation phenotype

The lateral line has not been previously implicated in initial inflation of the larval swim bladder. As such, it was unexpected to observe that lateral line mutants (*lhfpl5b^-/-^*) exhibit hyperinflated swim bladders by 5-6 dpf (Figure 1A). Based on our swim bladder volume and buoyancy measurements, we categorized each larva in the mutant (*n* = 31) and wild type (*n* = 33) populations as over-, regular-, or under-inflated (Figure 1B). On average, 53.7% (*SD* = 6.4%, *n* = 41) of lateral line mutants over-inflated their swim bladders and 7.3% (SD = 7.1%, *n* = 41) under-inflated. For wild type larvae (including *lhfpl5b^vo35^* heterozygotes), 0.0% (*SD* = 0.0%, *n* = 41) over-inflated and 4.9% (*SD* = 3.7%, *n* = 41) under-inflated their swim bladders during initial inflation on average (Figure 1C). The proportion of over-inflation was significantly different between mutant and wild type larvae (chi-square statistic: 70.45, *p* < 0.00001), while the proportion of under-inflation was not (chi-square statistic: 0.3546, *p* = 0.552). Average swim bladder volume of mutant fish was larger (*M* = 0.012 mm^3^, *SD* = 0.002 mm^3^) than wild type siblings (*M* = 0.0058 mm^3^, *SD* = 0.00043 mm^3^, *t*(80) = 17.4, *p* < 0.0001) (Figure 1D). The proportions of neutrally and positively buoyant larvae did not change when raised in total darkness (Supplemental Figure 1). This unexpected hyperinflation phenotype in *lhfpl5b^vo35^*homozygotes led us to investigate if the lateral line is contributing important sensory information during the surfacing behavior.

**Figure 1.**
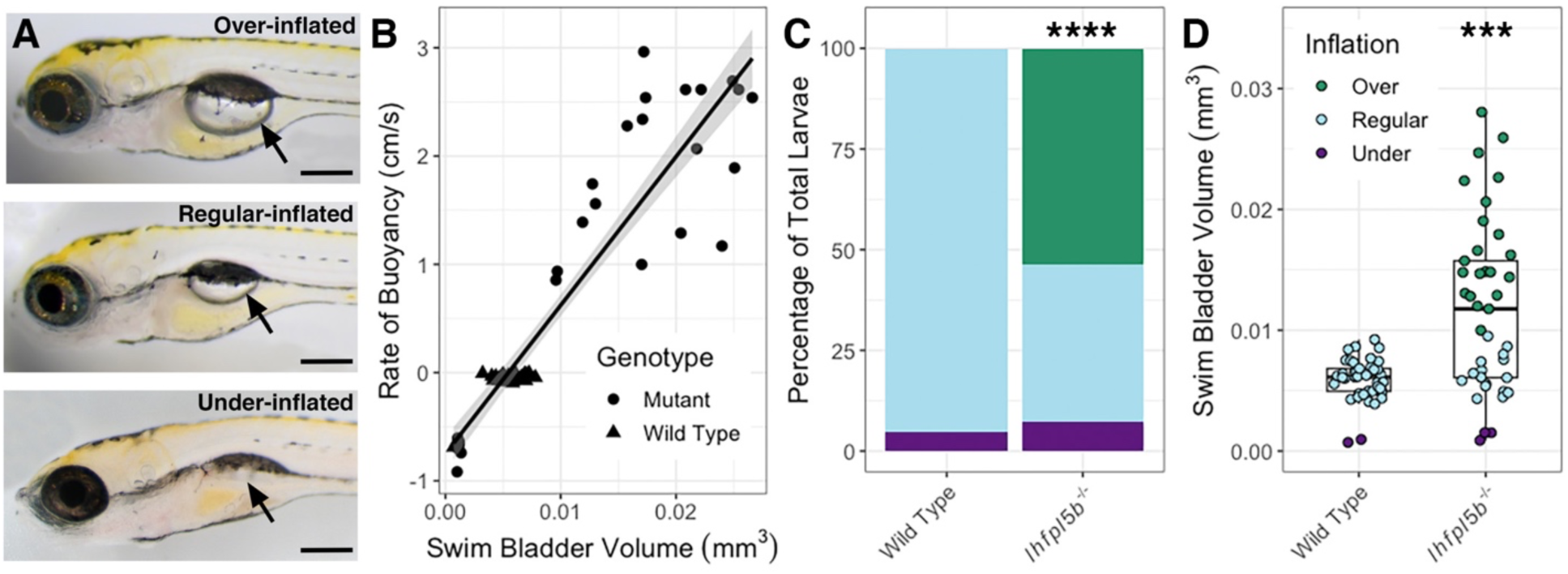
Hyperinflation of the swim bladder in lateral line mutants (*lhfpl5b*^-/-^). A) Representative examples of over-, regular, and under-inflated swim bladders (arrows) in larval zebrafish at 6 dpf (top: *lhfpl5b^-/-^*, others are wild type). B) Correlation between buoyancy and swim bladder volume. Line of best fit applied through the combination of mutant and wild type data (linear model, R-squared = 0.8574). C) Percentage of larvae that exhibit over-, regular, and under-inflation of the swim bladder, as defined by positive, neutral, and negative buoyancy. Legend from panel D applies to both panel C and D. A Chi-Squared Test was used to determine significance. D) Swim bladder volume (mm^3^) in wild type and *lhfpl5b* mutant larvae at 6 dpf. A Welch’s t-test was used to determine significance. *** = p < 0.001, **** = p < 0.0001. Scale bar = 0.2 mm.

### Access to the air-water interface is required for the hyperinflation phenotype of lateral line mutants

For most teleosts, initial swim bladder inflation requires an intake of exogenous air, typically from the water’s surface. To test if the hyperinflation phenotype of lateral line mutants requires access to surface air, we physically blocked access to the surface from 2-6 dpf (Figure 2A). As expected for wild type larvae, blocking access to the surface led to significantly more underinflated larvae (Figure 2B) and a significant decrease in the average swim bladder volume (Blocked: *M* = 0.0037 mm^3^, *SD* = 0.001 mm^3^, *n* = 49; Open: *M* = 0.0066 mm^3^, *SD* = 0.0009 mm^3^,

**Figure 2:**
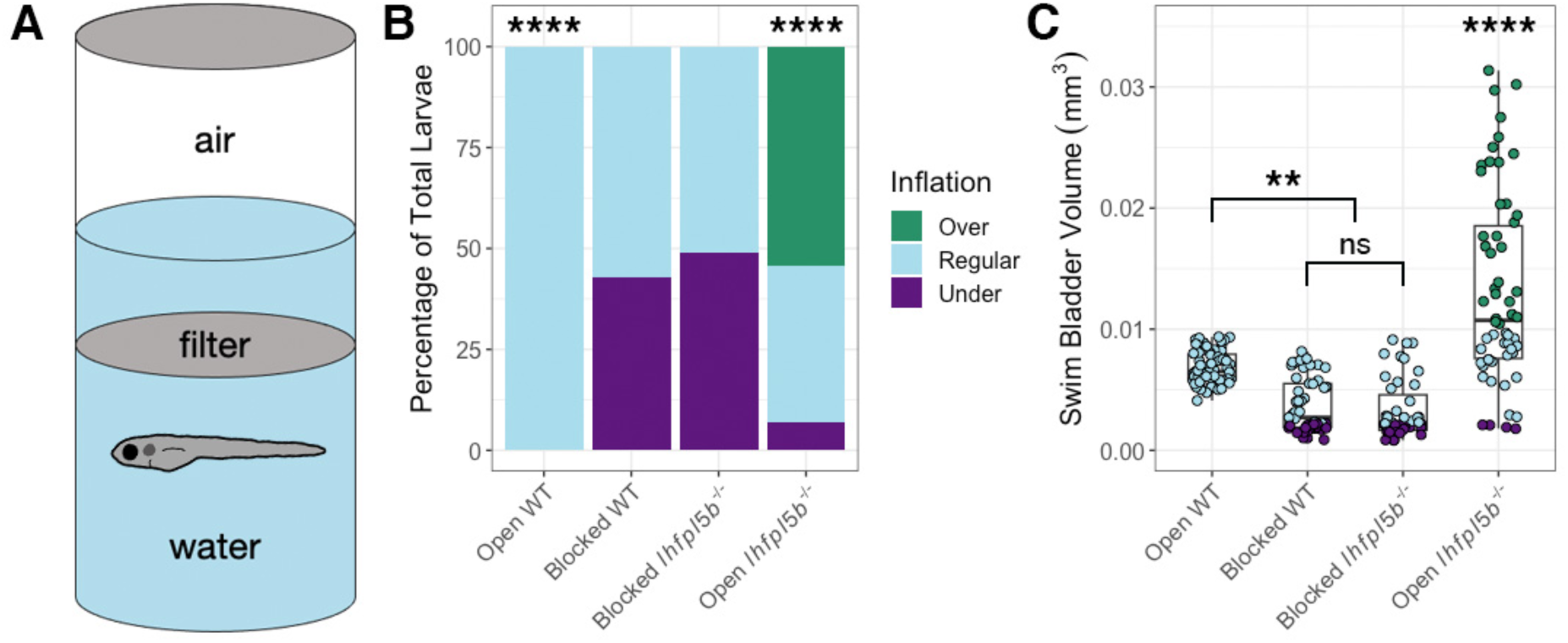
Blocking access to the surface eliminates the hyperinflation phenotype in lateral line mutants (*lhfpl5b*^-/-^). A) Diagram of experimental set up with a filter blocking access to the surface. B) Percentage of larvae that exhibit over-, regular, and under-inflation of the swim bladder at 6 dpf. A Chi-Squared Test was used to determine significance. C) Swim bladder volume (mm^3^) of larvae at 6 dpf. A One-Way ANOVA was used to determine significance. **** = p < 0.0001, ** = p < 0.01, ns = not significant. Full ANOVA table in supplemental tables.

*n* = 63) (One-way ANOVA with Tukey post-test, *p* = 0.002) (Figure 2C). For *lhfpl5b* mutant siblings, blocking access to the surface completely prevented the hyperinflation phenotype, with 0% (*SD* = 0.0%, *n* = 43) of blocked *lhfpl5b* mutants exhibiting positive buoyancy, which is significantly less than mutants with access to the surface (chi-square statistic: 70.45, *p* < 0.00001; Figure 2B). Likewise, the average swim bladder volume of blocked mutants (*M* = 0.0033 mm^3^, *SD* = 0.0015 mm^3^) was significantly smaller than mutants with open access (*M* = 0.014 mm^3^, *SD* = 0.0027 mm^3^, *n* = 58) (One-way ANOVA with Tukey post-test, *p* < 0.00001). Blocked mutant and wild type siblings had statistically similar swim bladder volumes (One-way ANOVA with Tukey post-test, *p* = 0.972). From these data, we concluded that the *lhfpl5b* mutant hyperinflation phenotype requires intake of exogenous air and is not caused by abnormal gas exchange from surrounding media nor excess internal gas production.

### Transgenic rescue of lateral line mutants restores normal inflation

We next examined if lateral line defects are responsible for the hyper-inflation phenotype of *lhfpl5b* mutants by restoring lateral line function in mutant larvae. We used the *Tg(myo6b:EGFP-lhfpl5a)vo23* transgene, which uses a hair cell specific promoter (*myo6b*) to drive a GFP-tagged *lhfpl5a* gene (Erickson et al., 2017). It has been shown that this *lhfpl5a* transgene rescues lateral line hair cell function in *lhfpl5b^-/-^ (or* lateral line) mutant fish (Erickson et al., 2020). Using this rescue method, we observed no cases of hyperinflation in transgenic mutants (0%, *SD* = 0.0%, *n* = 17), which was significantly different from non-transgenic mutants (chi-square statistic: 82.12, *p* < 0.00001; Figure 3A). The average volume of the transgenic mutant swim bladder (*M* = 0.0057 mm^3^, *SD* = 0.00001 mm^3^) was significantly smaller than non-transgenic, *lhfpl5b* mutant siblings (*M* = 0.0109 mm^3^, *SD* = 0.0031 mm^3^, *n* = 18) (One-way ANOVA with Tukey post-test, *p* = 0.000003), but statistically similar to that of wild type transgenics (*M* = 0.0057 mm^3^, *SD* = 0.0005 mm^3^, *n* = 57) (One-way ANOVA with Tukey post-test, *p* = 0.979, Figure 3B). Because the restoration of *lhfpl5* function specifically in hair cells mitigates hyperinflation in *lhfpl5b* mutants, we concluded that the phenotype is due to defects in sensory hair cell function and not caused by an unrecognized role for *lhfpl5b* in another sensory organ or the swim bladder itself.

**Figure 3:**
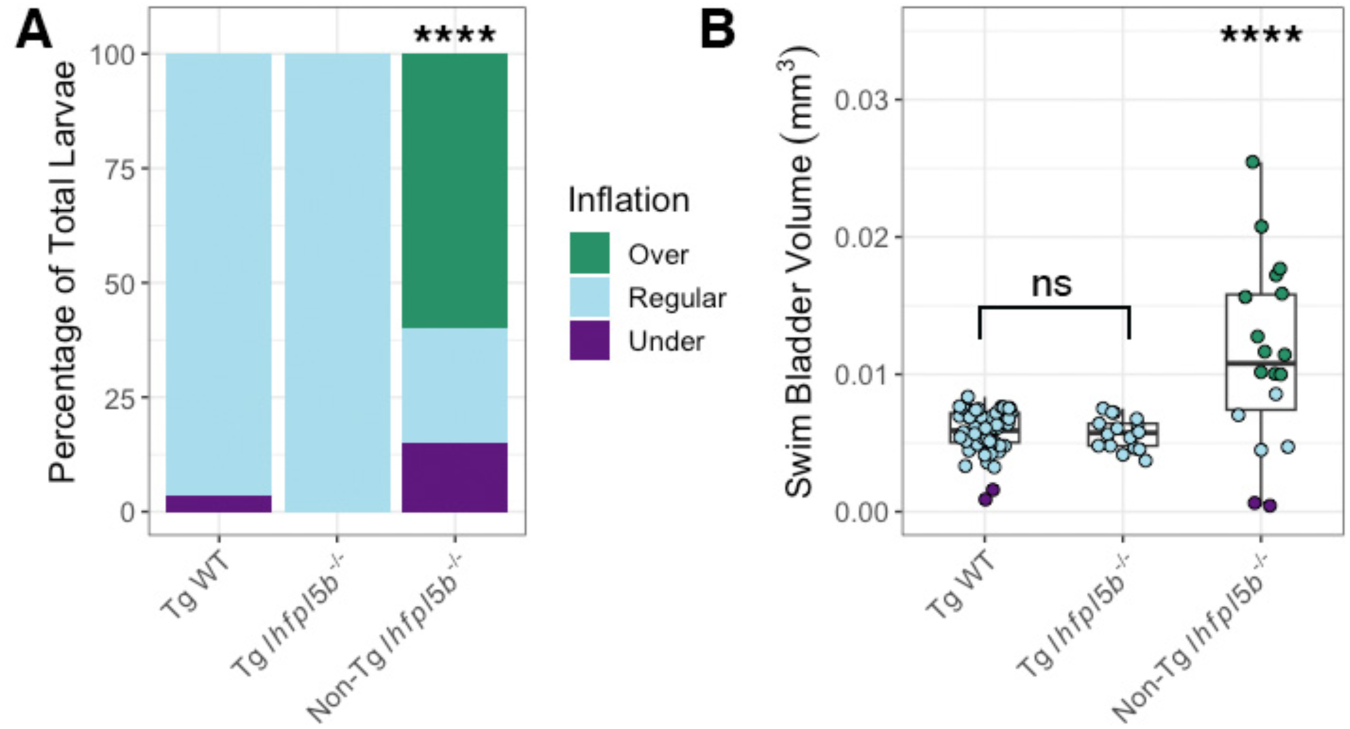
Rescue of lateral line mutants (*lhfpl5b*^-/-^) with the *Tg(myo6b:EGFP-lhfpl5a)vo23* transgene. A) Percentage of larvae that exhibit swim bladder under, regular, and over-inflation at 6 dpf. A Chi-Squared Test was used to determine significance. B) Swim bladder volume (mm^3^) of larvae at 6 dpf. A One-Way ANOVA was used to determine significance. **** = p < 0.0001, ns = not significant. Full ANOVA table in supplemental tables.

### Selective ablation of head neuromasts produces a hyperinflation phenotype in wild type larvae

To further test if lateral line defects are responsible for the hyper-inflation phenotype seen in *lhfpl5b* mutants, we ablated lateral line hair cells in wild type larvae using either neomycin sulfate (neo) or copper sulfate (CuSO4) (Supplemental Figure 2 and Figure 4). Administering either single or repeated neomycin treatments during the initial swim bladder inflation period of 3–4 dpf resulted in a significant increase in the proportion of positively buoyant larvae (single neo chi-square statistic: 9.95, *p* = 0.0016, *n* = 58; repeated neo chi-square statistic: 34.88, *p* < 0.0001, *n* = 47; Supplemental Figure 2A), but a non-significant increase in the average swim bladder volume (One-way ANOVA with Tukey post-test, single neo *p* = 1.0, repeated neo p = 0.578; Supplemental Figure 2B). By comparison, a single CuSO4 treatment resulted in a hyperinflation phenotype that was statistically indistinguishable from lateral line mutants. An average of 59.6% (*SD* = 7.7%, *n* = 72) of CuSO4-treated wild types over inflate their swim bladders, which was significantly different from wild type controls (chi-square statistic: 39.69, *p* < 0.0001, *n* = 44; Supplemental Figure 2A). The average swim bladder volume of the CuSO4-treated wild type larvae (*M* = 0.0126 mm^3^, *SD* = 0.0027 mm^3^) was statistically similar to the average swim bladder volume of untreated lateral line mutants (*M* = 0.0139 mm^3^, *SD* = 0.0007 mm^3^, *n* = 45) (One-way ANOVA with Tukey post-test, *p* = 0.813) (Supplemental Figure 2B).

**Figure 4:**
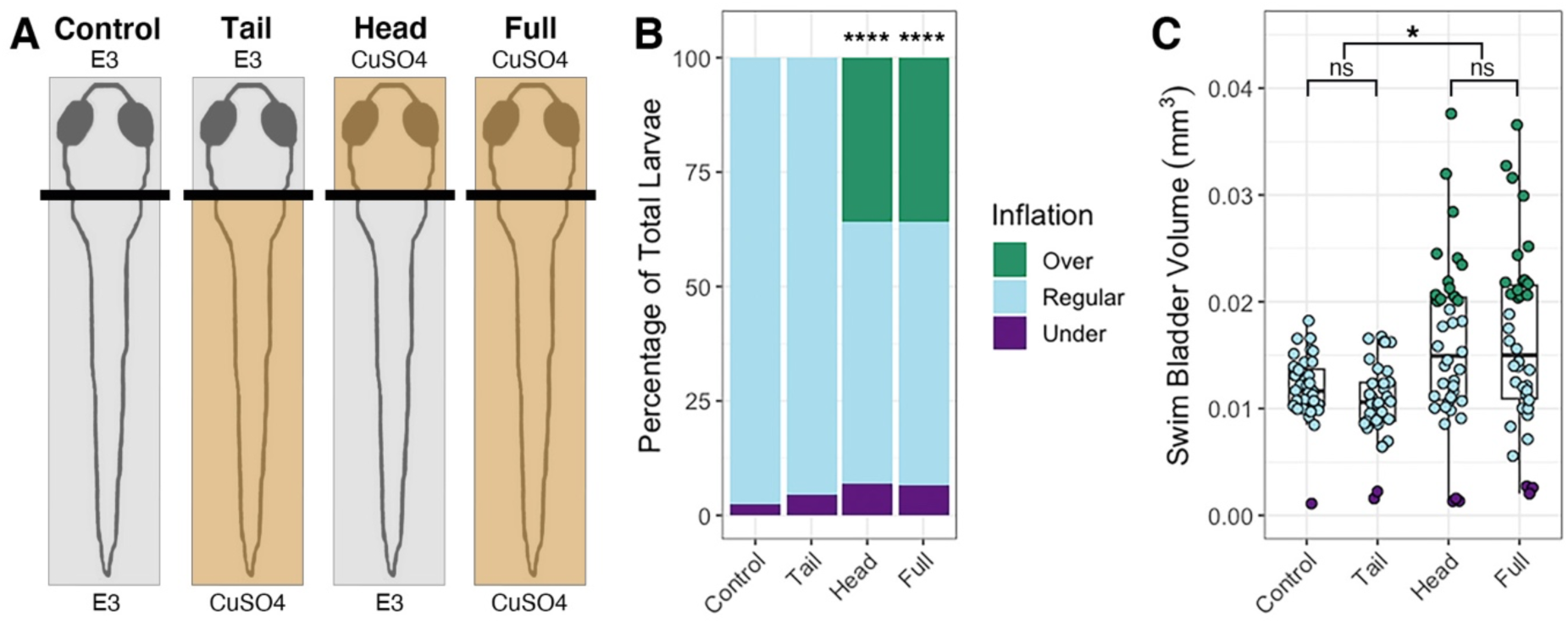
Head-specific ablations of the lateral line recapitulates the *lhfpl5b* mutant hyperinflation phenotype. A) Diagram of experiment. B) Percentage of larvae that exhibit swim bladder under, regular, and over-inflation at 5 dpf. A Chi-Squared Test was used to determine significance. C) Swim bladder volume (mm^3^) of larvae at 5 dpf. A One-Way ANOVA was used to determine significance. * = p < 0.05, **** = p < 0.0001, ns = not significant. Full ANOVA table in supplemental tables. Images of live larvae following treatment in Supplemental Figure 3.

Since the anterior lateral line is likely to be the only part of the sensory organ to interact with the air-water interface during surfacing, we predicted that disruption of the head neuromasts will result in hyperinflation and that trunk neuromasts will not play a role (Figure 4, Supplemental Figure 3). Selective ablation of the anterior region with CuSO4 resulted in an average of 35.9% (*SD* = 10.1%, *n* = 38) hyperinflated larvae, which was significantly different from unablated controls (chi-square statistic: 39.69, *p* < 0.0001; Figure 4B). Wild type larvae with head-specific ablations exhibited a significant increase in average swim bladder volume (*M* = 0.016 mm^3^, *SD* = 0.0013 mm^3^) compared to untreated wild type siblings (*M* = 0.0012 mm^3^, *SD* = 0.0013 mm^3^, *n* = 36) (One-way ANOVA with Tukey post-test, *p* = 0.0365) and statistically comparable swim bladder volumes to that of full CuSO4-treated larvae (*M* = 0.016 mm^3^, *SD* = 0.0029 mm^3^, *n* = 38) (One-way ANOVA with Tukey post-test, *p* = 0.987) (Figure 4C). Conversely, tail-specific CuSO4 treatments did not cause hyperinflation in wild type fish and resulted in statistically similar swim bladder volumes to untreated siblings (*M* = 0.012 mm^3^, *SD* = 0.0031 mm^3^, *n* = 34) (One-way ANOVA with Tukey post-test, *p* = 0.835) (Figure 4C). The *lhfpl5b* mutant hyperinflation phenotype was not affected by either neomycin or CuSO4 treatments (Supplemental Figure 2). Taken together, we concluded that the anterior region of the lateral line system is critical to surface detection during initial swim bladder inflation.

### Decreasing surface tension results in over-filling of the swim bladder

Surface tension is a characteristic that makes the air-water interface distinct from the rest of the water column. We predicted that larval fish sense this stimulus with their anterior lateral line neuromasts and that by decreasing the interfacial tension, we would observe abnormal swim bladder inflation in wild type larvae while lateral line mutants would be unaffected. The interfacial tension between oil and water is approximately half that of air and water (Johansen, 1924). We layered oil onto the surface of the embryo media and allowed larvae to perform surfacing behaviors between 2-6 dpf (Figure 5). Using this experimental design, we found that the average swim bladder volume of wild types exposed to an oil-water interface (*M* = 0.016 mm^3^, *SD* = 0.0029 mm^3^, *n* = 32) was significantly greater than the control group with access to an air-water interface (*M* = 0.007 mm^3^, *SD* = 0.001 mm^3^, *n* = 28) (One-way ANOVA with Tukey post-test, *p* < 0.0001). Furthermore, oil-exposed wild types had statistically similar swim bladder volumes to that of *lhfpl5b* mutants (*M* = 0.014 mm^3^, *SD* = 0.0038 mm^3^, *n* = 24) (One-way ANOVA with Tukey post-test, *p* = 0.886) (Figure 5C). Lateral line mutants overfilled their swim bladders to statistically similar proportions when presented with either an air-water or oil-water interface (*M* = 0.01 mm^3^, *SD* = 0.0026 mm^3^, *n* = 28) (One-way ANOVA with Tukey post-test, *p* = 0.133). We concluded that surface tension is one of the surface characteristics the lateral line is capable of detecting during initial swim bladder inflation.

**Figure 5:**
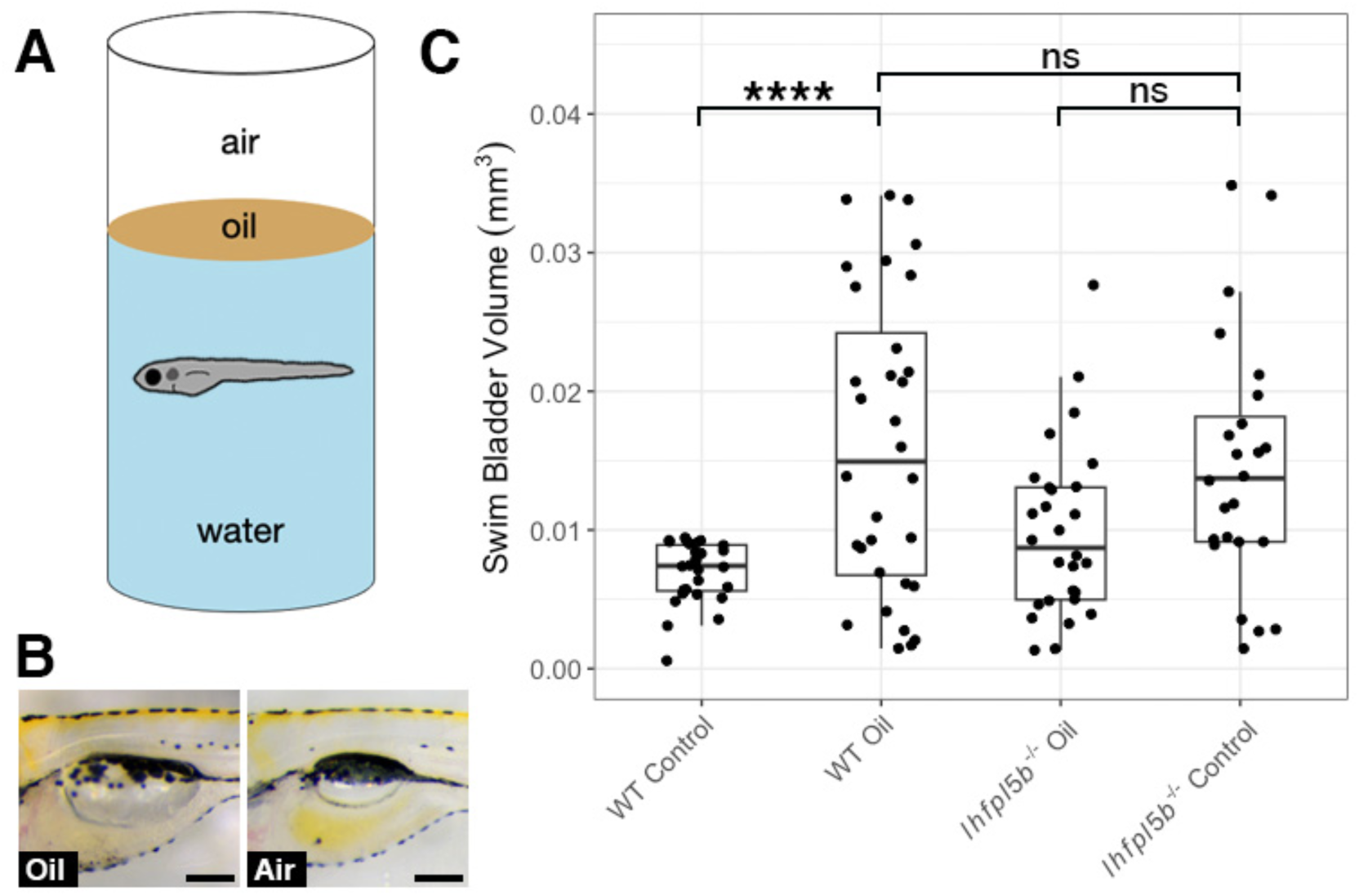
Decreasing interfacial surface tension with mineral oil results in over-filling of the swim bladder. A) Diagram of experiment with 2 mm layer of oil on surface. B) Images of swim bladder phenotypes resulting from surface oil and surface air exposure in wild type larvae at 6 dpf. C) Swim bladder volume (mm^3^) at 6 dpf. A One-Way ANOVA was used to determine significance. * = p < 0.05, ns = not significant. Scale bar = 0.1 mm. Full ANOVA table in supplemental tables.

### Optogenetic activation of lateral line hair cells prevents hyperinflation in lateral line mutant larvae

If hydrodynamic information is informing the surfacing behaviors of larval fish, then stimulus-independent manipulations of lateral line activity should alter their ability to correctly inflate their swim bladders. To test this, we created the *Tg(lhfpl5b2028:ChR2-EYFP-PA)* transgenic line that expresses the blue light activated channel, Channelrhodopsin-2, specifically in lateral line hair cells (See Methods, Supplemental Figure 4). We predicted that optogenetic stimulation of *lhfpl5b* mutants, whose lateral lines are insensitive to physical stimuli, would reduce the frequency of the hyperinflation phenotype. Consistent with these predictions, under blue-light stimulation, lateral line mutants with the *lhfpl5b:ChR2* transgene did not exhibit positive buoyancy and had a significant decrease in average swim bladder volume (*M* = 0.0046 mm^3^, *SD* = 0.0034 mm^3^) compared to unstimulated transgenic mutants (*M* = 0.012 mm^3^, *SD* = 0.0031 mm^3^) (One-way ANOVA with Tukey post-test, *p* = 0.00127) (Figure 6B). Transgenic wild types under blue light exhibited a non-significant decrease in swim bladder volume (*M* = 0.0052 mm^3^, *SD* = 0.004 mm^3^) compared to unstimulated transgenic siblings (*M* = 0.0079 mm^3^, *SD* = 0.002 mm^3^) (One-way ANOVA with Tukey post-test, *p* = 0.458). To demonstrate that blue light stimulation was reducing swim bladder volumes via activation of lateral line hair cells, we treated wild type *lhfpl5b:ChR2* larvae with CuSO4. As expected, we observed hyperinflation following CuSO4 treatment in stimulated wild type *lhfpl5b:ChR2* larvae, with an average swim bladder volume (WT: *M* = 0.012 mm^3^, *SD* = 0.0037 mm^3^) that was statistically similar to experimental non-transgenic *lhfpl5b* mutants (*M* = 0.011 mm^3^, *SD* = 0.0024 mm^3^) (One-way ANOVA with Tukey post-test, *p* = 0.998) (Supplemental Figure 4B). CuSO4 treatments in *lhfpl5b* mutants did not kill hair cells due to the non-functional MET channel, but rather acted as a control for potential behavioral side effects of the CuSO4 treatments. Overall, optogenetic activation of mechanically-insensitive lateral line organs in *lhfpl5b* mutants prevented the hyperinflation phenotype. Together, these data support the hypothesis that the lateral line provides crucial sensory information during initial swim bladder inflation in larval zebrafish.

**Figure 6:**
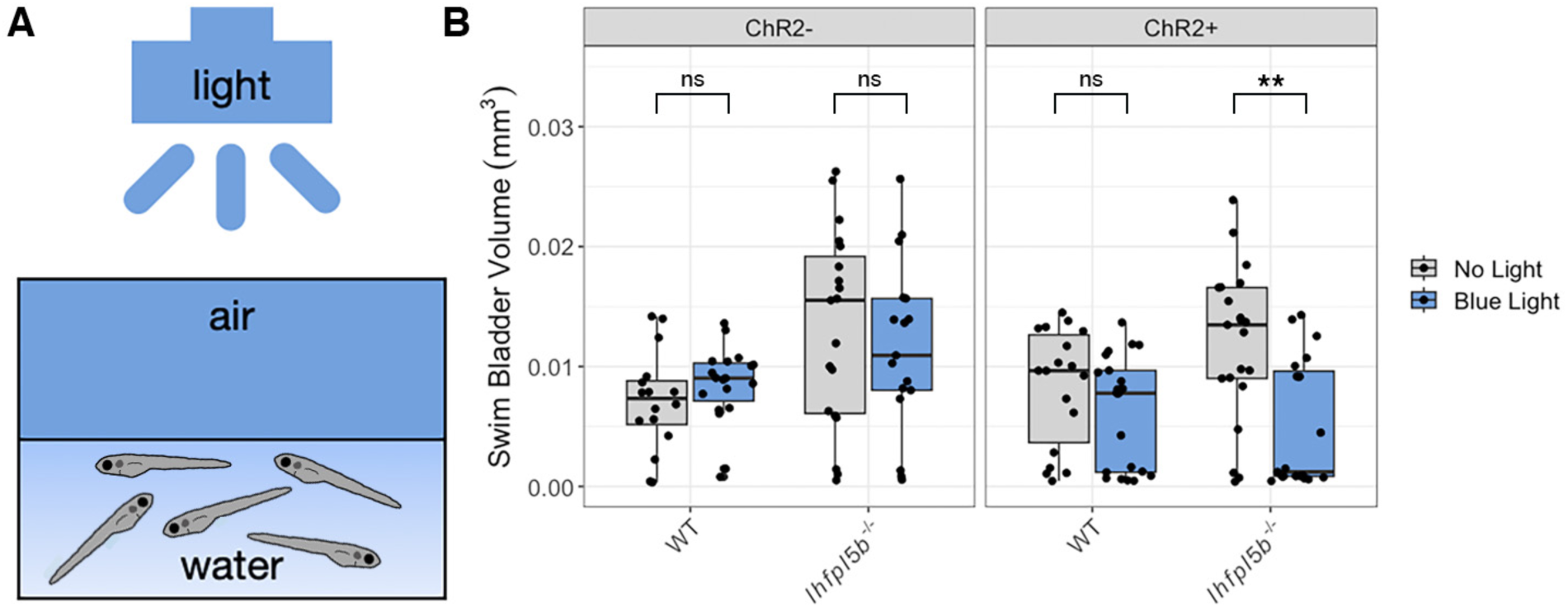
Optogenetic activation of lateral line hair cells with Channelrhodopsin-2 (ChR2) in wild type and *lhfpl5b* mutant larval zebrafish. A) Diagram of experiment with blue light stimuli delivered for 25 milliseconds at 1 second intervals. Treatments were started at 4 dpf and swim bladder volume was assessed at 6 dpf. B) Swim bladder volume (mm^3^) of larvae at 6 dpf, n = 16-21 per condition. One-Way ANOVA was used to determine significance. ** = p < 0.001, ns = not significant. Full ANOVA table in supplemental tables.

### Lateral line mutants exhibit abnormal surfacing behaviors

Our results suggested that larval fish are using their head neuromasts to sense the interfacial tension between air and water at the surface. Even in wild type zebrafish, the interaction with the surface during initial swim bladder inflation is not well characterized. To study this behavior in more detail, we quantified the number of unique visits to the surface and the amount of time per visit spent within the meniscus region from videos of wild type (n = 7) and mutant (n = 14) larvae at 4 dpf. These counts excluded larvae (n = 5 total: 1 wild type and 4 mutant) that under-inflated due to lack of surfacing events. *lhfpl5b* mutants that over-inflate their swim bladders made more visits (*M* = 87.3, *SD* = 17.4) to the surface compared to both wild type (*M* = 42.0, *SD* = 16.3) and mutant siblings that end up inflating to normal proportions (*M* = 31.9, *SD* = 19.2) (One-way ANOVA with Tukey post-test, *p* = 0.0004) (Figure 7A). Mutants destined for hyperinflation also spent more time (*M* = 139.0 s, *SD* = 31.6 s) at the surface than normally-inflated wild type (*M* = 51.9 s, *SD* = 24.7 s) and mutant siblings (*M* = 42.3 s, *SD* = 29.5 s) (One-way ANOVA with Tukey post-test, *p* = 0.00006) (Figure 7B). Since excess surfacing could be due to some mutants initiating the behavior earlier, we also documented the time of the first surfacing event and found no significant difference between neutrally buoyant and over-inflated larvae (Figure 7C). We concluded there is a correlation between how larvae interact with the surface during the initial swim bladder inflation period and their resulting swim bladder volume and buoyancy status.

**Figure 7:**
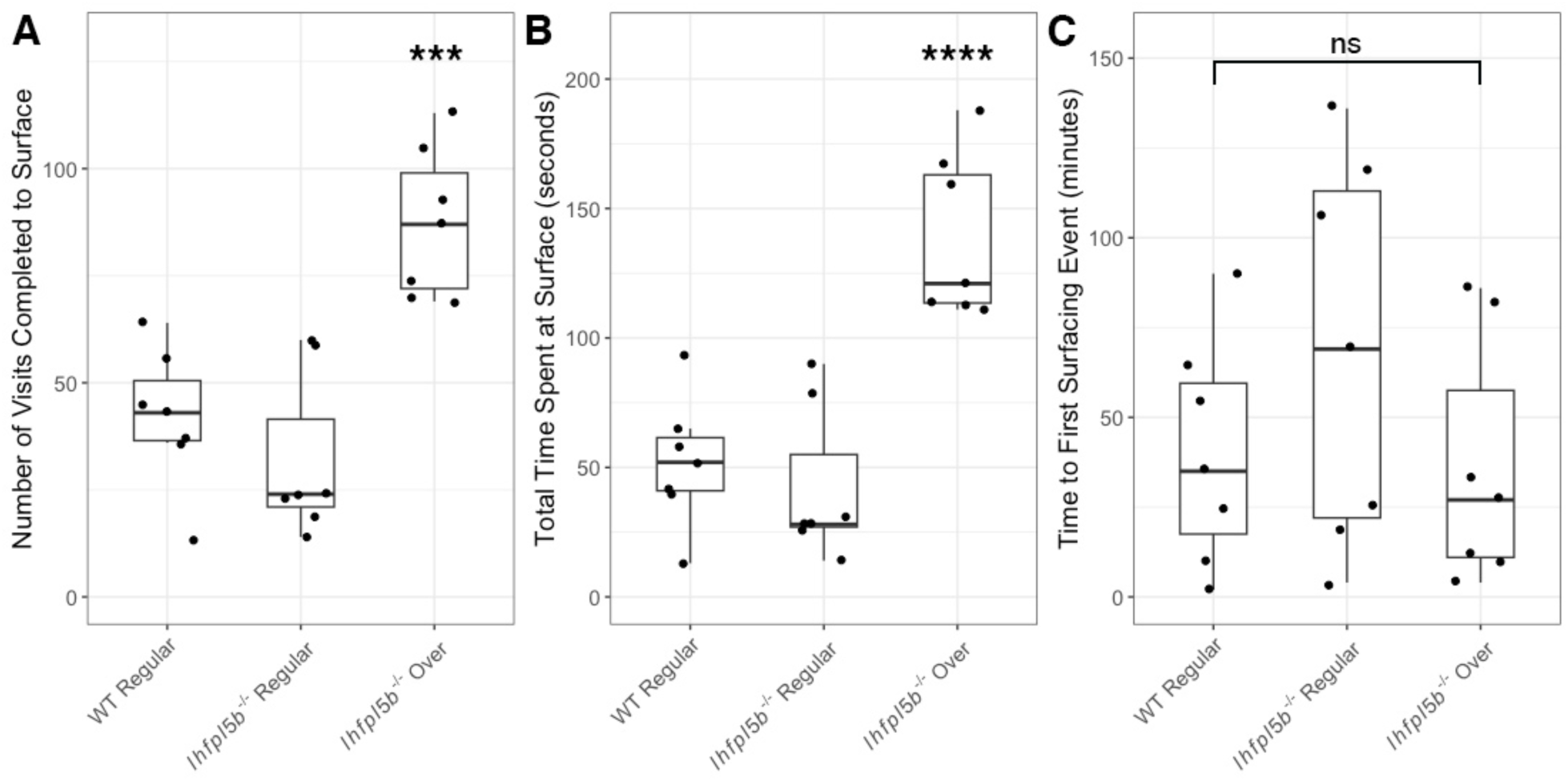
Quantification of the surfacing behaviors of wild type and *lhfpl5b* mutant zebrafish larvae during initial swim bladder inflation. A) Total number of individual visits taken by larval fish to the air-water interface on 4 dpf for 12 hours (9 AM – 9 PM). B) Total amount of time larval fish spend at the surface for all visits combined on 4 dpf. C) Time to first surface visit after access was allowed. A One-Way ANOVA was used to determine significance. *** = p < 0.001, ns = not significant.

## Discussion

The intake of exogenous air from the surface is a common feature for intitial swim bladder inflation in many species of fish (Furukawa et al., 2022; Goolish and Okutake, 1999; Jones and Marshall, 1953; Korsøen et al., 2012; v. Ledebur and Wunder, 1937). However, few studies have considered what sensory information is required to achieve neutral buoyancy. In this study, we describe a new role for the mechanosensory lateral line in mediating initial inflation of the swim bladder in zebrafish (Figure 8). Our data support a model in which larval zebrafish use their head neuromasts to detect the air-water interface in support of their transition to neutral buoyancy.

**Figure 8.**
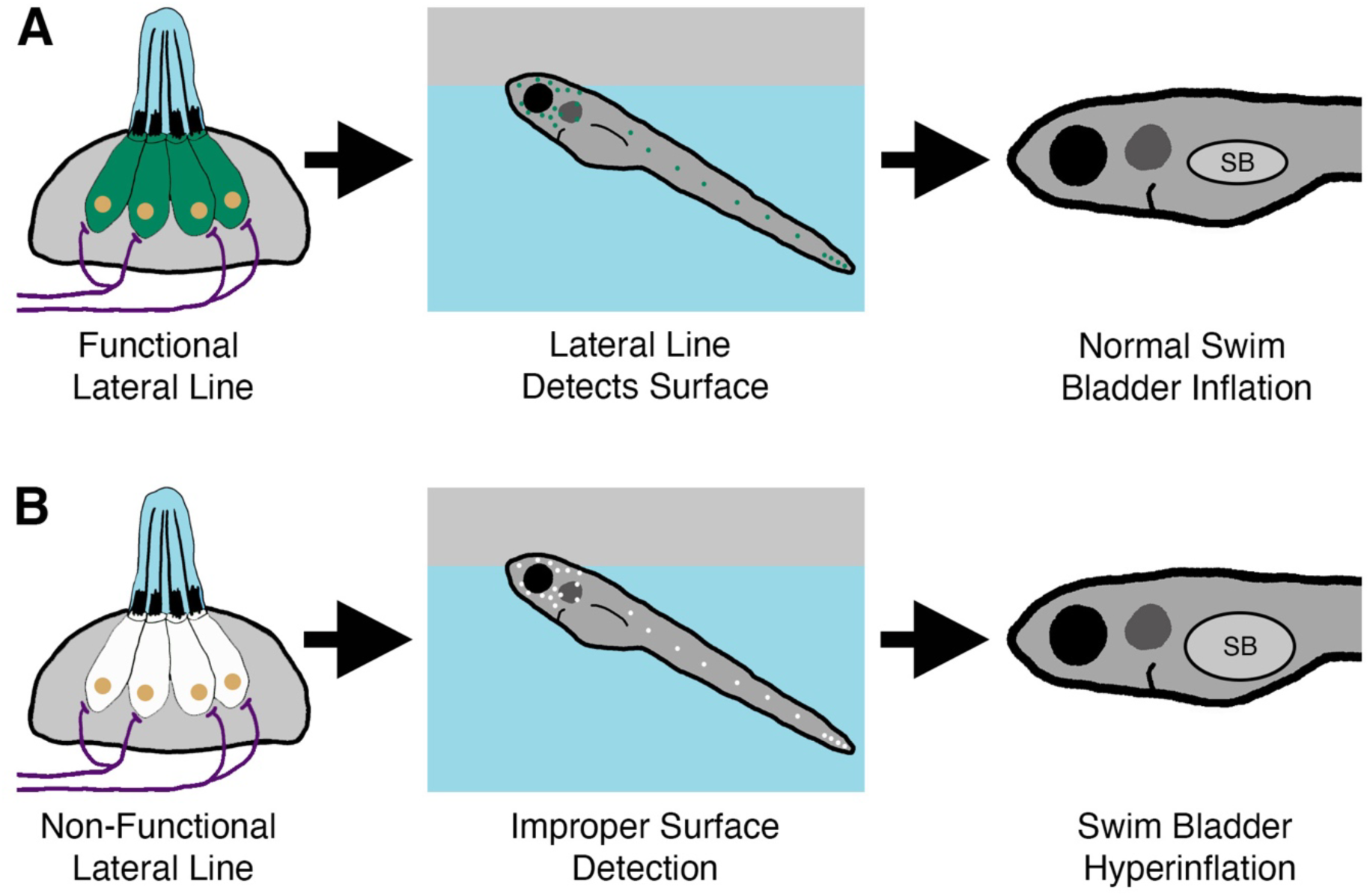
Summary of the surfacing behavior of wild type and lateral line-deficient zebrafish larvae. A) Wild type larvae use their lateral line hair cells (in green) to mediate interactions with the air-water interface during surfacing. As a result of accurate surface detection, larvae take in an appropriate volume of air for swim bladder (SB) inflation and achieve neutral buoyancy. B) Larvae with a genetic loss of lateral line function (hair cells in white) misinterpret the air-water interface, leading to increased interactions with the surface. Consequently, lateral line mutants *(lhfpl5b^-/-^)* take in an excess volume of air, resulting in hyperinflation of the swim bladder for approximately half of mutant larvae. Selective chemical ablations of the head neuromasts produce similar results, implicating the anterior lateral line as the primary sensory organ for interactions with the air-water interface during the surfacing behavior.

### Lateral line defects in lhfpl5b mutants are responsible for hyperinflation phenotypes

*lhfpl5b* mutant zebrafish are the first genetic model for the specific loss of lateral line function in any organism (Erickson et al., 2020). These “lateral line mutants” exhibit an uncommon swim bladder hyperinflation phenotype that prompted us to investigate this novel connection between lateral line sensation and initial inflation of the swim bladder (Figure 1). Previous studies have demonstrated that restricting access to surface air during the initial inflation period prevents swim bladder inflation in larval zebrafish (Goolish and Okutake, 1999; Lindsey et al., 2011). In addition, one study showed that some *solute carrier family 12 member 2* (*slc12a2 / nkcc1)* mutant zebrafish over-fill their swim bladders and that blocking access to the surface prevents this phenotype (Abbas and Whitfield, 2009). This gene is implicated in endolymph production in the inner ear, but further studies are required to determine the cause of over-inflation in *slc12a2* mutants. Likewise, we find that blocking access to the air-water interface eliminates over-inflation in lateral line mutants and greatly decreases average swim bladder volumes of both mutant and wild type larvae (Figure 2). Our method of blocking access to the air-water interface resulted in a small proportion of larvae (both wild type and mutant) inflating their swim bladder to normal proportions, though hyperinflation was never observed. Inflation was likely achieved by larvae accessing air bubbles that remain in the water after submerging the filter (Goolish and Okutake, 1999). Overall, the hyperinflation phenotype in lateral line mutants requires an intake of exogenous air, pointing to a sensory defect in *lhfpl5b* mutants rather than a physiological one.

In support of this sensory-deficit hypothesis, we show the hyperinflation phenotype is prevented by either transgenic rescue of *lhfpl5* function specifically in hair cells or optogenetic stimulation of mechanically-insensitive lateral line hair cells in *lhfpl5b* mutants (Figures 3 and 6). Furthermore, chemical ablation of lateral line hair cells in wild type larvae prior to initial inflation can phenocopy the *lhfpl5b* mutant. A single treatment with CuSO4 caused similar swim bladder over-inflation proportions to lateral line mutants (Supplemental Figure 2). Repeated neomycin treatments over 3-4 dpf caused the next highest percentage of swim bladder over-inflation and a single neomycin treatment only resulted in a few cases of wild type over-inflation. These results agree with previous work showing rapid hair cell regeneration following neomycin treatment and comparatively slower recovery (approximately 24 hours for full recovery) after CuSO4 exposure (Hernández et al., 2006; Mackenzie and Raible, 2012; Murakami et al., 2003; Santos et al., 2006; Venuto and Erickson, 2021). Thus, via multiple genetic and chemical approaches, our data show that lateral line hair cell activity can modulate swim bladder inflation in larval zebrafish.

Ototoxins have been extensively used to study lateral line-mediated behaviors in free-swimming larvae and adult fish (Baker and Montgomery, 1999; Bleckmann, 2006; Coffin and Ramcharitar, 2016; Montgomery et al., 1997; Pavlov and Tyuryukov, 1993). However, we are not aware of any behavioral studies where the lateral line was disrupted prior to larvae achieving neutral buoyancy, likely explaining why this connection between the lateral line and initial swim bladder inflation has not been noted in current literature. Interestingly, an 1896 study predicted a relationship between the lateral line and swim bladder inflation, since disruption of the lateral line in adult goldfish resulted in “swelling of the swim bladder” and positive buoyancy (Richard, 1896). However, it is not clear if the phenotypic similarities between these adult goldfish and our experimental larval zebrafish share the same underlying causes. Regarding roles for the lateral line in early life behaviors, a close parallel to our findings is the report that unhatched red-eyed treefrogs use their lateral line organ to mediate mechanosensory-cued hatching (Jung et al., 2020). In this case, the frog lateral line can sense vibrational stimuli, such as that produced by predators, and promote escape hatching behaviors even before the onset of vestibular function. Similarly, our data suggest that the lateral line is modulating early locomotor activities in zebrafish prior to the development of fully formed vestibular or rheotactic responses (Bagnall and Schoppik, 2018; Mo et al., 2010; Olive et al., 2016). As such, the involvement of lateral line in the surfacing behavior may represent the earliest behavioral role for the lateral line yet described.

### Head neuromasts are required for normal swim bladder inflation, likely by sensing surface tension

We expanded on the full-body lateral line ablation experiments with specific ablations of either the head or truck neuromasts with CuSO4. Previous studies have shown that head neuromasts are involved with detection of surface features and prey capture (Bleckmann, 1988; Carrillo and McHenry, 2016; Müller and Schwartz, 1982; Schwartz and Hasler, 1966). Indeed, ablation of the head neuromasts in wild type larvae yielded similar swim bladder hyperinflation levels to larvae that experienced full body treatments. While this specific experiment did not include *lhfpl5b* mutants, we find that head-treated larvae generally recapitulated the hyperinflation phenotype of *lhfpl5b* mutants. Ablation of the tail neuromasts had no effect on either swim bladder volume or buoyancy (Figure 4). These findings suggest that the head neuromasts interact with the air-water interface during surfacing and that the posterior lateral line is dispensible for achieving neutral buoyancy. As such, our findings also provide additional support for the anterior lateral line as a central mediator of surface-related behaviors in fish.

The interfacial tension between air and water represents a unique hydrodynamic stimulus that may be detected by the lateral line. The tension between oil and water is half that of air and water (Johansen, 1924). Thus, by adding a layer of oil to the surface, we predicted that surface detection would become increasingly difficult and result in over-filling of the swim bladder. We find that, when given access to an oil-water interface, wild type larvae over-fill their swim bladders with oil to a statistically similar volume as lateral line mutants who overinflate with air (Figure 6). We interpret this result as support for the hypothesis that the lateral line senses interfacial tension between air and water as larvae interact with the surface during initial swim bladder inflation. This interpretation is further supported by the observation that *lhfpl5b* mutants (who are already insensitive to interfacial tension) exhibit similar swim bladder volumes when exposed to either oil or air. However, a possible complementary interpretation for these data is that larvae who fill their swim bladders with oil may be deprived of the normal feedback on buoyancy levels as they intake from the surface. Since oil is denser than air, oil-filled larvae are negatively buoyant. In this scenario, larvae may continue to visit the surface for air intake in an attempt to achieve neutral buoyancy. However, future studies are required to examine if such a feedback mechanism exists in larval zebrafish.

### Swim bladder over-inflation is correlated with excess time spent at the air-water interface

Since our data suggest that the air-water interface is providing a distinct stimulus to the lateral line, we investigated whether surface interactions differed between hyperinflated *lhfpl5b* mutants and their wild type counterparts. As predicted, over-inflated mutants took more visits *and* spent more time at the surface than neutrally-buoyant mutant and wild type siblings (Figure 7). Excess breaching of the surface occurred despite over-inflated and neutrally buoyant larvae exhibiting similar timing for their first surfacing attempts. While it is unclear whether larvae ingest air during the entirety of the duration spent above the air-water interface, it is known that surface breaches are observed during initial inflation (Goolish and Okutake, 1999; Lindsey et al., 2010; Rieger and Summerfelt, 1998). Therefore, the increase in surface visits likely leads to an increase in air deposition to the swim bladder in lateral line mutants.

### Multisensory integration during the surfacing behavior

Interestingly, approximately half of all lateral line mutants inflate their swim bladders to normal proportions and continue to develop into viable adults. To account for this fact, we hypothesize that additional somatosensory, photosensory, or mechanosensory systems may compensate for the loss of lateral line sensation and help mutant larvae achieve neutral buoyancy (Riley and Moorman, 2000; Suchocki and Sepulveda-Villet, 2019; Trotter et al., 2003). The lateral line is involved several multi-sensory behaviors, including flow orientation (Bak-Coleman et al., 2013; Coombs et al., 2020; Liao, 2006), prey capture (Carrillo and McHenry, 2016; Nelson et al., 2002; New, 2002), and predator avoidance (Free et al., 2019; Jung et al., 2020; Stewart et al., 2013), and we predict that surfacing can be added to this list.

The photosensory and mechanosensory systems exhibit distinct organizational patterns for primary input to the central nervous system — even the anterior and posterior lateral line ganglia primary projections are somatotopically organized in zebrafish larvae (Alexandre et al., 1999; Fame et al., 2006; Hale et al., 2016; Liao and Haehnel, 2012; McCormick, 1989; Mirjany and Faber, 2011). The Mauthner neurons integrate multisensory input at the level of the hindbrain, but the surfacing behavior is unlikely to use this circuit. For higher order processing however, information from the lateral line, vestibular, and photosensory systems are integrated in various brain regions. The thalamus is one such sensory integration center in zebrafish (Mueller, 2012). Socially-relevant mechanosensory and photosensory information is represented in a thalamic region marked by the expression of *parathyroid hormone 2* (*pth2*) and *somatostatin 7* (*sst7*) (Anneser et al., 2020; Kappel et al., 2022; Venuto et al., 2022). Vestibular information is also represented in the thalamus (Favre-Bulle et al., 2018; Migault et al., 2018), though it is not clear if the exact same region is involved. Photosensory, lateral line, and vestibular/auditory information is also integrated in the tectum (Favre-Bulle et al., 2018; Filosa et al., 2016; Isa et al., 2021; Migault et al., 2018). In fact, neurons in the periventricular layer (PVL) of the tectum, which is similar to the superior colliculus in mammals, respond to visual, auditory, and lateral line inputs (Ingle, 1973; King et al., 1996; Perrault et al., 2011; Thompson et al., 2016). However, it is hypothesized that most of these modalities are processed in parallel at younger stages of development (Thompson et al., 2016). It is worth noting that these sensory systems are in developing stages during surfacing behaviors.

Photosensory cues did not alter the buoyancy status of either wild type or *lhfpl5b* mutant larvae (Supplemental Figure 1), suggesting that, at most, photosensory systems play a subtle modulatory role in the surfacing behavior of larval zebrafish. The vestibular system is likely required to coordinate the vertical orientation and climbing portions of surfacing (Bagnall and Schoppik, 2018; Ehrlich and Schoppik, 2017; Ehrlich and Schoppik, 2019). Meanwhile, our data support the idea that the lateral line is responsible for the surface detection and breaching aspects of surfacing. However, given that both the vestibular and lateral line systems are providing spatial information in during the surfacing behavior, there may be crosstalk between them. For example, the vestibular system must quickly adapt to a dynamic increase in buoyancy over the course of multiple breaching events, so feedback from the lateral line during surface interactions may help fine tune the vestibular sense. However, the link between these systems have not been well-investigated during deliberate behaviors like surfacing (Chagnaud et al., 2017; Liu et al., 2021). So, while sensory integration exists in zebrafish and seems necessary for larvae to achieve neutral buoyancy, the integration dynamics are unclear during the intial swim bladder inflation timeframe and require future investigation.

## Supporting information

Venuto Supplemental ANOVA tables v4

## Acknowledgements

The authors would like to thank Dr. Fadi Issa (ECU) for his support and technical advice. The authors also thank the Issa, Trapani, and Erickson lab members for fish care and maintenance.

## Author contributions

Conceptualization: A.V., T.E.; Methodology: A.V., J.G.T., T.E.; Validation: A.V.; Formal analysis: A.V.; Investigation: A.V.; Resources: S.T-B., J.G.T; Writing–original draft: A.V., T.E.; Writing–review & editing: A.V., J.G.T., T.E..; Visualization: A.V.; Supervision: T.E.; Project administration: T.E.; Funding acquisition: J.G.T., T.E.

---

**Supplemental Figure 1.**
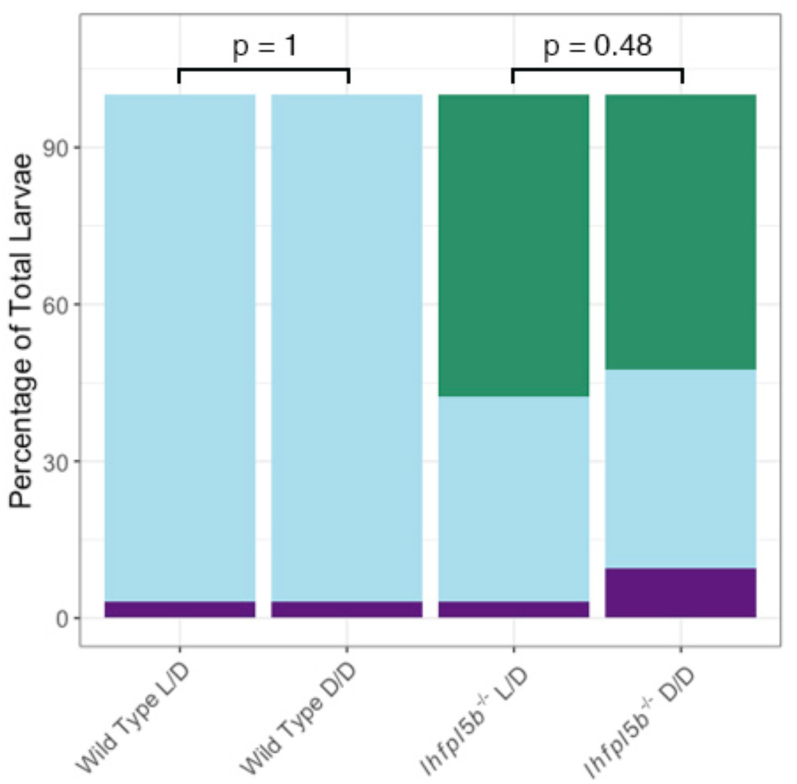
Buoyancy proportions of larvae raised in total darkness. Percentage of larvae that exhibit swim bladder under, regular, and over-inflation raised in either a 14:10 light/dark cycle (L/D) or total darkness (D/D). A Chi-Squared Test was used to assess significance. Both Wild Type L/D and Wild Type D/D over-inflation proportions: *M* = 0.0%, *SD* = 0.0%, *n* = 32. *lhfpl5b^-/-^* L/D over-inflation proportion: *M* = 57.5%, *SD* = 6.6%, *n* = 30. *lhfpl5b^-/-^* D/D over-inflation proportion: *M* = 52.4%, *SD* = 4.1%, *n* = 30.

**Supplemental Figure 2.**
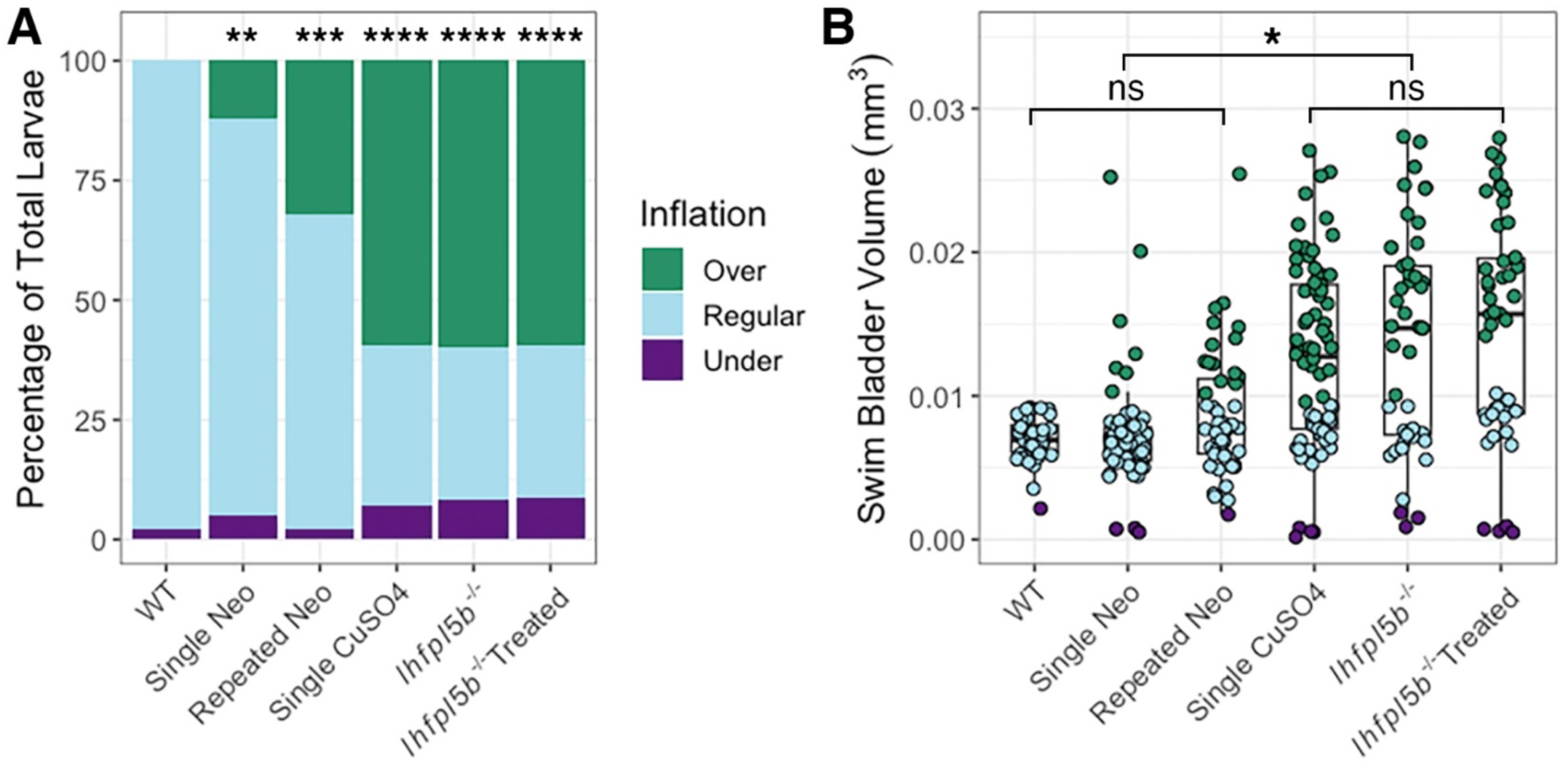
Ototoxic ablations of the lateral line. A) Percentage of larvae that exhibit swim bladder under, regular, and over-inflation. A Chi-Squared Test was used to assess significance. B) Swim bladder volume (mm^3^) in 5 dpf larvae. A One-Way ANOVA was used to determine significance. **** = *p* < 0.0001, *** = *p* < 0.001, ** = *p* < 0.01, ns = no significance. A full ANOVA table can be found in the supplemental tables.

**Supplemental Figure 3.**
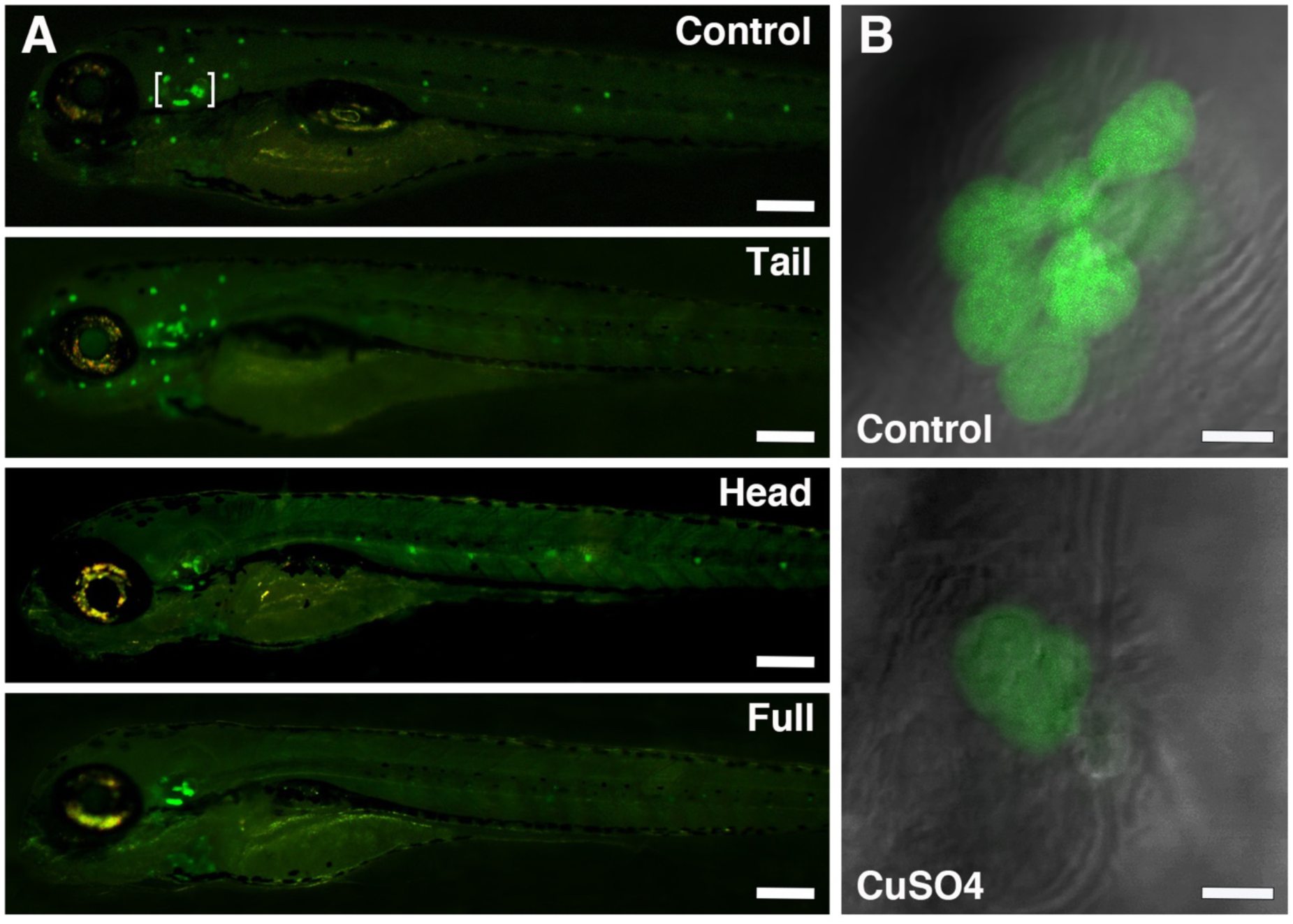
Anterior and posterior-specific lateral line ablations with CuSO4. A) Representative images of live *Tg(myo6b:eGFP-pA)vo68* transgenics at 4 dpf, two hours after CuSO4 treatment. This transgene labels hair cells of the lateral line and inner ear (indicated by white brackets on the Control image), and treatment only affects lateral line hair cells so inner ear expression remains post-treatment. B) Representative images from untreated Control and CuSO4-treated L1 neuromasts.

**Supplemental Figure 4:**
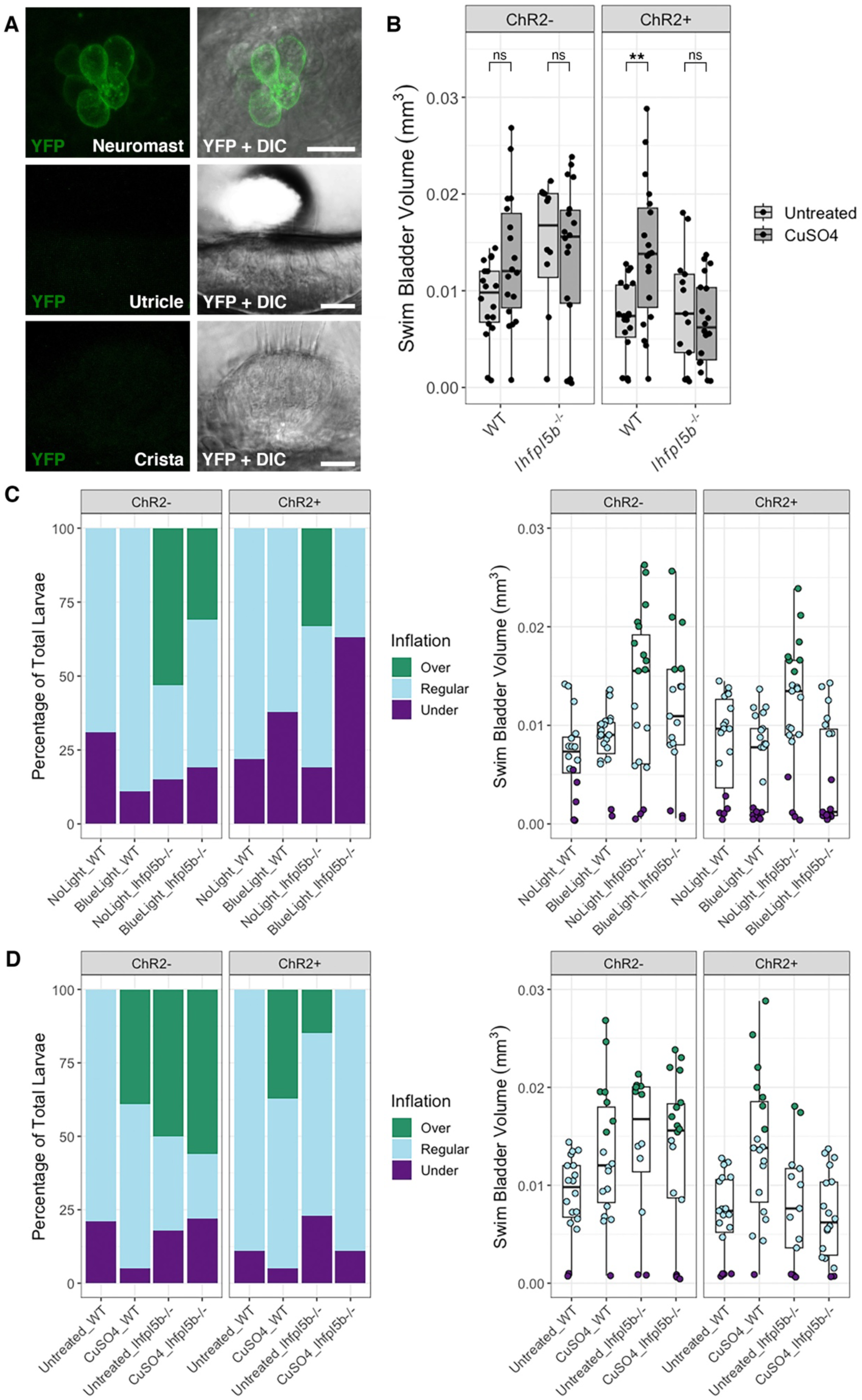
Channelrhodopsin-2 (ChR2) expression in *Tg(lhfpl5b2028:ChR2-EYFP-pA)* transgenics. A) Images of ChR2-YFP in hair cells in a 2 dpf neuromast, utricle, and crista where fluorescence expression is only seen in the neuromasts of the lateral line. First panel is YFP only (false-colored green) and the second panel is a merge of YFP and DIC. Scale bars = 10 uM. B) Swim bladder volume (mm^3^) of 6 dpf larvae from the experimental condition (blue light overhead). A One-Way ANOVA was used to determine significance. ** = *p* < 0.00724, ns = no significance. A full ANOVA table can be found in the supplemental tables. C) Percentages and box plots of experiment in Figure 6 to show inflation categories. D) Percentages and box plots of experiment in Supplemental Figure 4B to show inflation categories.

