## Supplementary material for "A sensation for inflation: initial swim bladder inflation in larval zebrafish is mediated by the mechanosensory lateral line": Venuto Supplemental ANOVA tables v4

Venuto et al. 2023 ANOVA tables

**Figure 2** - Blocking Surface Access - ANOVA Table (type II tests)

| Effect | DFn | DFd | F | p | p<0.05 | ges |
|---|---|---|---|---|---|---|
| Status | 3 | 209 | 55.021 | 2.96E-26 | * | 0.441 |
| group1 | group2 | estimate | conf.low | conf.high | p.adj | p.adj.signif |
| Blocked MUT | Blocked WT | 0.000407 | -0.00202 | 0.00283 | 0.972 | ns |
| Blocked MUT | Open MUT | 0.00984 | 0.00751 | 0.0122 | 9.33E-15 | **** |
| Blocked MUT | Open WT | 0.0035 | 0.00121 | 0.0058 | 6.00E-04 | *** |
| Blocked WT | Open MUT | 0.00943 | 0.00718 | 0.0117 | 1.24E-14 | **** |
| Blocked WT | Open WT | 0.0031 | 0.000888 | 0.0053 | 0.002 | ** |
| Open MUT | Open WT | -0.00634 | -0.00845 | -0.00423 | 1.98E-12 | **** |

**Figure 3** - Transgenic Rescue - ANOVA Table (type II tests)

| Effect | DFn | DFd | F | p | p<0.05 | ges |
|---|---|---|---|---|---|---|
| Status | 2 | 89 | 21.699 | 2.11E-08 | * | 0.328 |
| group1 | group2 | estimate | conf.low | conf.high | p.adj | p.adj.signif |
| Tg MUT | Tg WT | 0.000173 | -0.00194 | 0.00228 | 0.979 | ns |
| Non-Tg MUT | Tg MUT | 0.00567 | -0.00825 | -0.00309 | 3.20E-06 | **** |
| Non-Tg MUT | Tg WT | 0.0055 | -0.00756 | -0.00344 | 2.60E-08 | **** |

**Figure 4** – Region Specific Lateral Line Ablations - ANOVA Table (type II tests)

| Effect | DFn | DFd | F | p | p<0.05 | ges |
|---|---|---|---|---|---|---|
| Status | 3 | 142 | 7.442 | 1.15E-03 | * | 0.136 |
| group1 | group2 | estimate | conf.low | conf.high | p.adj | p.adj.signif |
| Head | Control | -0.00398 | -0.00778 | -0.000175 | 0.0365 | * |
| Head | Full | 0.000479 | -0.00327 | 0.00423 | 0.987 | ns |
| Head | Tail | -0.00524 | -0.0091 | -0.00138 | 0.0031 | ** |
| Control | Full | 0.00446 | -0.000655 | 0.00826 | 0.0145 | * |
| Control | Tail | -0.00126 | -0.00517 | 0.00264 | 0.835 | ns |
| Full | Tail | -0.00572 | -0.00958 | -0.00186 | 9.90E-04 | *** |

**Figure 5** – Surface Oil - ANOVA Table (type II tests)

| Effect | DFn | DFd | F | p | p<0.05 | ges |
|---|---|---|---|---|---|---|
| Status | 3 | 108 | 8.428 | 4.40E-05 | * | 0.19 |
| group1 | group2 | estimate | conf.low | conf.high | p.adj | p.adj.signif |
| MUT Control | MUT Oil | -0.00398 | -0.00477 | 0.000914 | 0.133 | ns |
| MUT Control | WT Control | 0.000479 | -0.00763 | -0.00194 | 0.0037 | ** |
| MUT Control | WT Oil | -0.00524 | 0.00154 | 0.00705 | 0.886 | ns |
| MUT Oil | WT Control | 0.00446 | -0.00286 | 0.0026 | 0.0523 | ns |
| MUT Oil | WT Oil | -0.00126 | 0.0063 | 0.0116 | 0.0125 | * |
| WT Control | WT Oil | -0.00572 | 0.00916 | 0.0144 | 9.10E-05 | **** |

**Figure 6** – Channelrhodopsin-2 - ANOVA Table (type II tests)

| Effect | DFn | DFd | F | p | p<0.05 | ges |
|---|---|---|---|---|---|---|
| Status | 7 | 142 | 4.734 | 8.40E-05 | * | 0.189 |
| group1 | group2 | estimate | conf.low | conf.high | p.adj | p.adj.signif |
| Con.Neg.Mut. | Con.Neg.WT. | -0.00582 | -0.0113 | -0.000303 | 0.035 | * |
| Con.Neg.Mut. | Exp.Neg.Mut. | 0.00145 | -0.00688 | 0.00397 | 0.895 | ns |
| Con.Neg.Mut. | Exp.Neg.WT. | -0.00472 | -0.00999 | 0.000555 | 0.0955 | ns |
| Con.Neg.WT. | Exp.Neg.Mut. | 0.00437 | -0.0013 | 0.01 | 0.187 | ns |
| Con.Neg.WT. | Exp.Neg.WT. | 0.0011 | -0.00442 | 0.00662 | 0.953 | ns |
| Exp.Neg.Mut. | Exp.Neg.WT. | -0.00327 | -0.00869 | 0.00216 | 0.394 | ns |
| Con.Pos.Mut. | Con.Pos.WT. | -0.00298 | -0.0077 | 0.00175 | 0.354 | ns |
| Con.Pos.Mut. | Exp.Pos.Mut. | -0.00687 | -0.0115 | -0.00221 | 0.00127 | ** |
| Con.Pos.Mut. | Exp.Pos.WT. | -0.00563 | -0.0102 | -0.00109 | 0.00893 | ** |
| Con.Pos.WT. | Exp.Pos.Mut. | -0.00389 | -0.00873 | 0.000946 | 0.158 | ns |
| Con.Pos.WT. | Exp.Pos.WT. | -0.00265 | -0.00737 | 0.00207 | 0.458 | ns |
| Exp.Pos.Mut. | Exp.Pos.WT. | 0.00124 | -0.00342 | 0.00589 | 0.897 | ns |

**Figure 7** – Behavior Surface Visits - ANOVA Table (type II tests)

| Effect | DFn | DFd | F | p | p<0.05 | ges |
|---|---|---|---|---|---|---|
| Status | 2 | 18 | 19.513 | 3.11E-05 | * | 0.684 |
| group1 | group2 | estimate | conf.low | conf.high | p.adj | p.adj.signif |
| MUT Over | MUT Regular | -55.4 | -79.5 | -31.3 | 4.20E-05 | **** |
| MUT Over | WT Regular | -45.3 | -69.4 | -21.2 | 4.10E-04 | *** |
| MUT Regular | WT Regular | 10.1 | -14 | 34.3 | 0.542 | ns |

**Figure 7** – Behavior Surface Time - ANOVA Table (type II tests)

| Effect | DFn | DFd | F | p | p<0.05 | ges |
|---|---|---|---|---|---|---|
| Status | 2 | 18 | 24.063 | 8.21E-06 | * | 0.728 |
| group1 | group2 | estimate | conf.low | conf.high | p.adj | p.adj.signif |
| MUT Over | MUT Regular | -96.7 | -136 | -57.5 | 1.70E-05 | **** |
| MUT Over | WT Regular | -87.1 | -126 | -47.9 | 6.30E-05 | **** |
| MUT Regular | WT Regular | 9.57 | -29.6 | 48.8 | 0.81 | ns |

**Figure 7** – Time to First Visit - ANOVA Table (type II tests)

| Effect | DFn | DFd | F | p | p<0.05 | ges |
|---|---|---|---|---|---|---|
| Status | 2 | 18 | 1.295 | 0.298 | - | 0.126 |
| group1 | group2 | estimate | conf.low | conf.high | p.adj | p.adj.signif |
| MUT Over | MUT Regular | 32.1 | -23.5 | 87.8 | 0.326 | ns |
| MUT Over | WT Regular | 3.86 | -51.8 | 59.5 | 0.983 | ns |
| MUT Regular | WT Regular | -28.3 | -83.9 | 27.4 | 0.415 | ns |

**Supplemental Figure 2** – Lateral Line Ablations - ANOVA Table (type II tests)

| Effect | DFn | DFd | F | p | p<0.05 | ges |
|---|---|---|---|---|---|---|
| Status | 5 | 306 | 18.314 | 6.63E-16 | * | 0.23 |
| group1 | group2 | estimate | conf.low | conf.high | p.adj | p.adj.signif |
| Mutant | Mutant Treat | 0.00089 | -0.00253 | 0.00431 | 0.976 | ns |
| Mutant | Repeat Neo | -0.00509 | -0.00849 | -0.00168 | 3.51E-04 | * |
| Mutant | Single CuSO4 | -0.00109 | -0.00419 | 0.00201 | 0.915 | ns |
| Mutant | Single Neo | -0.00662 | -0.00986 | -0.00337 | 1.88E-07 | **** |
| Mutant | Wild Type | -0.00682 | -0.0103 | -0.00336 | 5.40E-07 | **** |
| Mutant Treat | Repeat Neo | -0.00598 | -0.00936 | -0.00259 | 1.06E-05 | **** |
| Mutant Treat | Single CuSO4 | -0.00198 | -0.00506 | 0.0011 | 0.441 | ns |
| Mutant Treat | Single Neo | -0.00751 | -0.0107 | -0.00428 | 1.71E-09 | **** |
| Mutant Treat | Wild Type | -0.00771 | -0.0112 | -0.00427 | 7.61E-09 | **** |
| Repeat Neo | Single CuSO4 | 0.004 | 0.000936 | 0.00706 | 2.94E-03 | ** |
| Repeat Neo | Single Neo | -0.00153 | -0.00473 | 0.00168 | 0.746 | ns |
| Repeat Neo | Wild Type | -0.00174 | -0.00516 | 0.00169 | 0.694 | ns |
| Single CuSO4 | Single Neo | -0.00553 | -0.00841 | -0.00265 | 1.18E-06 | **** |
| Single CuSO4 | Wild Type | -0.00573 | -0.00886 | -0.00261 | 3.98E-06 | **** |
| Single Neo | Wild Type | -0.000206 | -0.00347 | 0.00306 | 1 | ns |

**Supplemental Figure 4** – Channelrhodopsin-2 Ablations - ANOVA Table (type II tests)

| Effect | DFn | DFd | F | p | p<0.05 | ges |
|---|---|---|---|---|---|---|
| Status | 7 | 127 | 4.678 | 1.08E-04 | * | 0.205 |
| group1 | group2 | estimate | conf.low | conf.high | p.adj | p.adj.signif |
| Tr.Neg.MUT. | Tr.Neg.WT. | -0.000109 | -0.00605 | 0.00583 | 1 | ns |
| Tr.Neg.MUT. | Un.Neg.MUT. | 0.000809 | -0.00583 | 0.00745 | 0.988 | ns |
| Tr.Neg.MUT. | Un.Neg.WT. | -0.00431 | -0.0102 | 0.00163 | 0.232 | ns |
| Tr.Neg.WT. | Un.Neg.MUT. | 0.000918 | -0.00572 | 0.00756 | 0.983 | ns |
| Tr.Neg.WT. | Un.Neg.WT. | -0.0042 | -0.0101 | 0.00174 | 0.253 | ns |
| Un.Neg.MUT. | Un.Neg.WT. | -0.00512 | -0.0118 | 0.00152 | 0.187 | ns |
| Tr.Pos.MUT. | Tr.Pos.WT. | 0.00702 | 0.0022 | 0.0118 | 0.00158 | ** |
| Tr.Pos.MUT. | Un.Pos.MUT. | 0.00128 | -0.00406 | 0.00661 | 0.922 | ns |
| Tr.Pos.MUT. | Un.Pos.WT. | -0.000246 | -0.00457 | 0.00507 | 0.999 | ns |
| Tr.Pos.WT. | Un.Pos.MUT. | -0.00574 | -0.011 | -0.000466 | 0.0277 | * |
| Tr.Pos.WT. | Un.Pos.WT. | -0.00677 | -0.0115 | -0.00202 | 0.00207 | ** |
| Un.Pos.MUT. | Un.Pos.WT. | -0.00103 | -0.00631 | 0.00425 | 0.955 | ns |
